# *MCF2* is linked to a complex perisylvian syndrome and affects cortical lamination

**DOI:** 10.1101/733444

**Authors:** Aude Molinard-Chenu, Joël Fluss, Sacha Laurent, Michel Guipponi, Alexandre G Dayer

## Abstract

The combination of congenital bilateral perisylvian syndrome (CBPS) with lower motor neuron dysfunction is unusual and suggests a potential common genetic insult affecting basic neurodevelopmental processes. Here we identify a putatively pathogenic missense mutation in the *MCF2* gene in a boy with CBPS. Using *in utero* electroporation to genetically manipulate cortical neurons during corticogenesis, we demonstrate that the mouse *Mcf2* gene controls the embryonic migration of cortical projection neurons. Strikingly, we find that the CBPS-associated *MCF2* mutation impairs cortical laminar positioning, supporting the hypothesis that alterations in the process of embryonic neuronal migration can lead to rare cases of CBPS.

## Introduction

Congenital bilateral perisylvian syndrome (CBPS) is a rare malformation of the cerebral cortex, most often associated with the following clinical signs: pseudobulbar palsy with oromotor apraxia, mild bilateral spastic palsy, cognitive impairment and epilepsy^1^. Despite this well-reported clinical phenotype, there is often a diagnostic delay, especially when some of the clinical features are missing in the early stages of the disorder. CBPS is considered by certain authors as a peculiar form of cerebral palsy affecting predominantly bulbar muscles, but some patients also display congenital contractures as clinical and neurophysiological signs consistent with an involvement of the lower motor neuron (MN) ^2–4^. The mechanism underlying this unusual combination of both central and peripheral motor impairment is unknown, but it most probably involves a common genetic insult affecting basic cellular processes regulating the development of MNs and cortical projection neurons (PNs). During the embryonic period, PNs and MNs are generated in germinal ventricular zones then migrate radially to reach their final anatomical position and interestingly, they share common molecular regulators ^5–7^. Here, we identify a missense mutation in the MCF.2 Cell Line Derived Transforming Sequence (*MCF2)* gene in a boy with CBPS associated to lower MN dysfunction. Using functional *in vivo* studies in mice, we find that this CBPS-associated human mutation has a pathogenic effect on the process of embryonic PNs migration *in vivo*.

## Patient, materials and methods

### Clinical description

The study was approved by the local ethics committee and parental consent was obtained for the patient who is a 9 year-old boy from Lebanon, born premature at 31-weeks gestation after a prolonged oligoamnios. Both parents are healthy and not related. There are no siblings. At birth, the patient exhibited bilateral clubfeet and hip dislocations, and an asymmetric paraparesis leading to the diagnosis of congenital arthrogryposis. During childhood, a severe developmental speech and language impairment was observed along with excessive drooling and feeding problems. Clinical examination of the patient confirmed a pseudobulbar palsy, an asymmetric lower limb distal paresis with no movement below the knees and severe muscle wasting. Right knee jerk was present but left knee jerk was absent, as were ankle jerks bilaterally.

### MRI study

All MRI data were acquired with a dedicated head coil on a 1.5T machine (Avanto Siemens Erlangen Germany).

### Exome sequencing

Exome of the patient was captured using the Agilent SureSelet QXT Human All Exon V5 kit and sequenced on a HiSeq2000 instrument (Illumina). Reads mapping and variant calling were performed using BWA 0.7.13, Picard 2.9.0, GATK HaplotypeCaller 3.7 and annotated with annovar 2017-07-17 and UCSC RefSeq (refGene) downloaded on 2018-08-10. The variants were searched in various databases including dbSNP151, gnomAD 2.1, ClinVar 2018 and HGMD 2016. Pathogenicity prediction scores were obtained for missense variants using SIFT, PolyPhen2, MutationTaster (MT), CADD.

### Mouse in utero electroporation

Animal experiments were conducted according to the Swiss and international guidelines, and approved by the local animal care committee. Embryos from time pregnant embryonic day (E)14.5 CD1 mice were electroporated in the lateral VZ of the dorsal pallium as described previously^8^. All constitutive expression of shRNAs and cDNAs was driven by the human U6 promoter, in the PLKO.1 vector for the shRNAs and pUB6/V5-His A (pUB6) vector for h*MCF2* (kind gift from Danny Manor), TOM and GFP. The following shRNAs were electroporated in equal ratios in control and experimental conditions: *Mcf2* shRNA (TRCN0000042653, Thermoscientific), and *Scramble* shRNA (mature sense: CCTAAGGTTAAGTCGCCCTCG, Addgene). The G4A missense mutation was induced in h*MCF2* plasmid using site-directed mutagenesis (InFusion Kit, Takara).

## Results

### MRI and nerve conduction studies

MRI scan of the spinal cord revealed a homogeneous thin spinal cord without the physiological lumbar enlargement (Fig 1.A). MRI scans of the brain revealed bilateral asymmetric perisylvian polymicrogyria extending to the parietal cortex on the right side. The sylvian sulci also displayed an abnormal configuration bilaterally (Fig 1.B-B’’).

**Figure 1:**
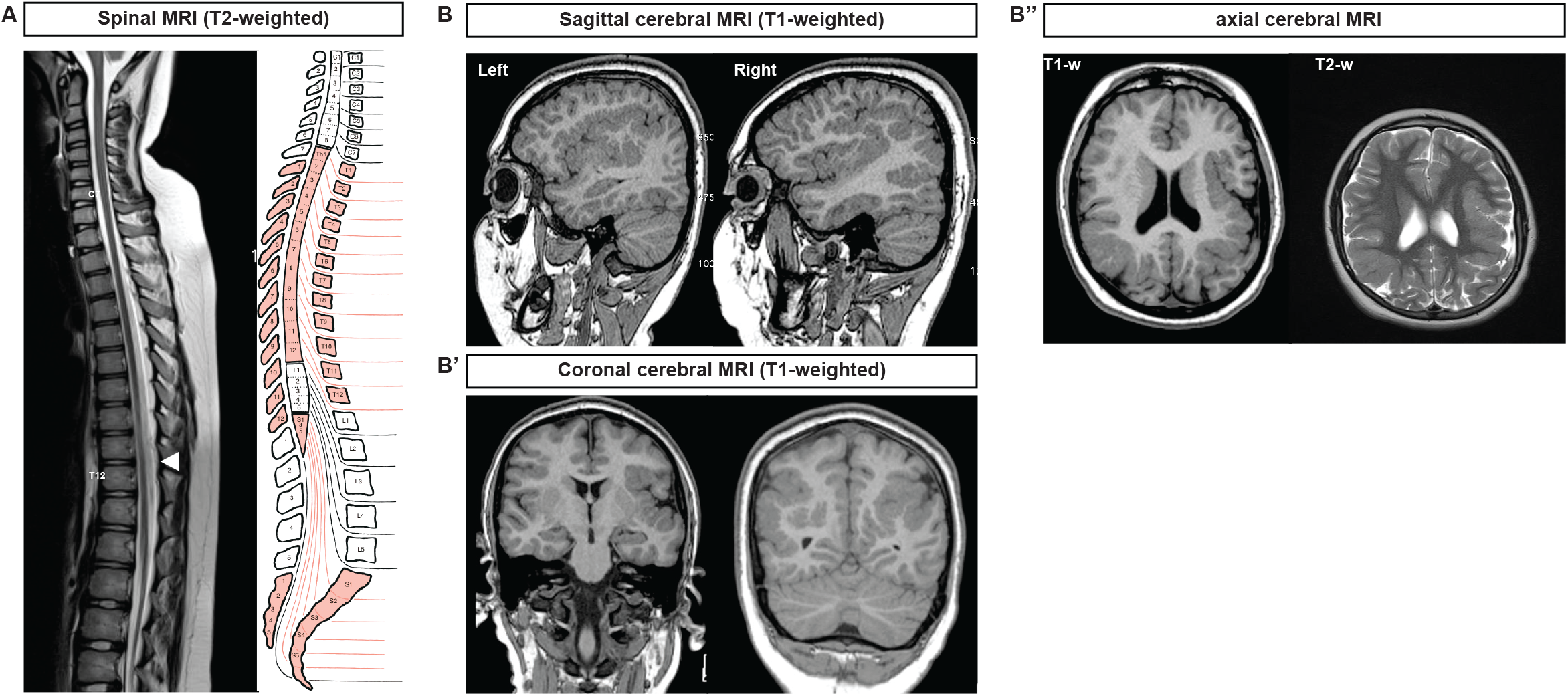
MRI study shows a thin spinal cord and bilateral perisylvian polymicrogyria. (A) Spinal cord T2-weighted(w) MRI shows a thin spinal cord from the dorsal spinal cord to the conus medullaris, without the physiological lumbar enlargement (arrow). (B-B’’) Cerebral T1-w and T2-w MRI display bilateral asymmetric perisylvian polymicrogyria extending to the parietal cortex on the right side. The sylvian sulci also display an abnormal configuration bilaterally.

Additionally, nerve conduction studies demonstrated a pure motor impairment from L4-S2 on both sides typical of a lower MN dysfunction.

### Exome sequencing

Exome sequencing of the proband and his healthy parents revealed a putatively damaging missense mutation inherited from his mother in the *MCF2* gene, located on chromosome Xq27, (NM_005369.5): c.4G>A, p.(Ala2Thr) in exon 1. This variant is absent from the gnomAD database (https://gnomad.broadinstitute.org/) and is predicted as pathogenic by all used algorithms except Polyphen2 which classified it as likely pathogenic. *MCF2*, also known as *DBL*, is a member of a large family of guanine nucleotide exchange factors (GEFs) that modulates the activity of Rho GTPases^9,10^. Given that GEFs have been shown to act as key regulators of cellular migration^11^, we aimed to test whether the CBPS-associated *MCF2* mutation affected cortical PNs *in vivo*.

### In vivo mouse studies

*Mcf2* was found to be expressed in control E17.5 PNs in a previously published RNA sequencing dataset^8^ and its subcellular cytoplasmic expression in cortical PNs at postnatal day (P)0 was visualized following overexpression of HA-tagged MCF2 by *in utero* electroporation of PNs progenitors at E14.5 in the dorsal pallium (Fig 2.A). A short hairpin RNA (shRNA) targeting *Mcf2* induced a 40% knock-down (KD) of the mRNA expression of *Mcf2*, as demonstrated by qRT-PCR of HEK 293T cells transfected with pUB6-*mMcf2* and control *Scram* or *Mcf2* targeting shRNA plasmids (Fig 2.B). Using *in utero* electroporation to genetically manipulate migrating PNs in mice, we next determined whether developmental KD of *Mcf2* regulated the migration of PNs. Strikingly, the laminar positioning of PNs was altered by *Mcf2* KD. Indeed, ectopic PNs were found in the lower layers of the somatosensory cortex in comparison to control (*Scram*) PNs, which were mostly located in superficial cortical layers. Interestingly, this phenotype was fully rescued by the overexpression of the WT human ortholog of *Mcf2* (*hMCF2*), resistant to the effect of the mouse-targeted shRNA, proving the specificity of the *Mcf2-*KD-related migratory defect. In contrast, overexpression of the CBPS-associated mutated *MCF2* failed to rescue the migratory deficit induced by *Mcf2* shRNA, indicating a pathogenic effect of the missense MCF2 mutation (*G4A - p.A2T*) on cortical PNs migration (Figure 2.C).

**Figure 2:**
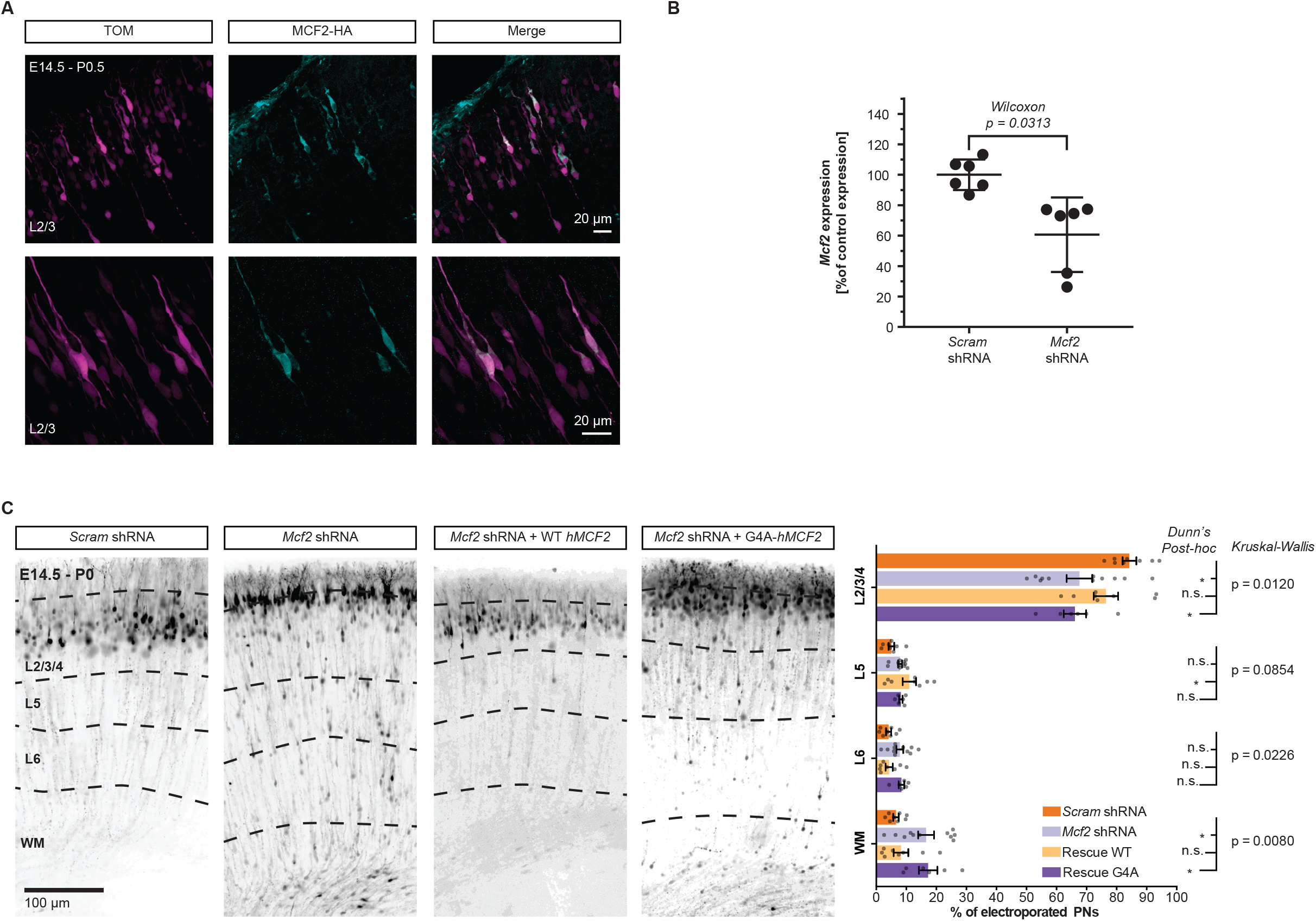
Mouse in vivo study: Mcf2 knock-down (KD) alters the laminar positioning of projection neurons (PNs) at P0.5 and the G4A missense mutation alters the migratory function of MCF2. (A) The overexpression of h*MCF2* by E14.5 electroporation in S1 PNs displays a cytoplasmic expression at P0.5. (B) q-RT-PCR of mRNA extracts from HEK-293T cells transfected with *mMcf2* and *Mcf2* or *Scramble* (*Scram*) shRNA shows a 40% knock-down (KD) of the mRNA expression by *Mcf2* shRNA, compared to *Scram* shRNA. Error bars = 95% C.I. (C) shRNA-mediated *Mcf2* KD by *in utero* electroporation at E14.5 dramatically impairs the laminar positioning of PNs in S1 at P0.5. This phenotype is fully rescued by the overexpression of the shRNA-resistant hMCF2 but the congenital bilateral perisylvian syndrome (CBPS)-associated missense mutation G4A prevents any rescue. n = 6-11 brains per condition from ≥ 3 separate litters. Error bars = 95% C.I.

## Discussion

In this short report, we have attempted to explore the underlying genetic cause of an usual but repetitively reported combination of perisylvian polymicrogyria and lower motor neuron dysfunction. These findings have led us to consider a genetic insult affecting basic cellular processes common to the maturation of both PNs and MNs. We have been able to identify an X-linked recessive inherited CBPS-associated missense mutation in *MCF2* (G4A - p.A2T). Using cell-type genetic manipulation in mouse embryos, we observed that *Mcf2*-KD impairs corticogenesis by altering the laminar positioning of PNs *in vivo*. Strikingly, the *MCF2* (G4A-p.A2T) mutation was found to have pathogenic effects on neuronal migration, suggesting that this developmental process could be altered in rare cases of CBPS.

The cell-specificity of the migratory deficit described in PNs remains to be explored in MNs, but core biological pathways involved in the migration of PNs appear to be shared with MNs^5^. In particular, MCF2 is part of the DBL family of RhoGEFs that represents critical regulators of cellular migration^11^. RhoGEFs have been involved in regulating neuronal migration through the REELIN pathway^12,13^ and the SEMAPHORIN/PLEXIN pathway^14,15^. The molecular underpinnings of the pathological effect of the missense mutation in *MCF2* (G4A - p.A2T) are beyond the scope of the present study, but based on the structure of MCF2, we hypothesize that it could impact the function of the SEC14 domain of the protein, thus affecting its subcellular localization and its GEF activity^10^. Interestingly, another missense mutation in *MCF2* has been associated with schizophrenia^16^, a complex neurodevelopmental disorder involving risk-genes regulating neuronal migration^8^. Additionally, one missense mutation in *MCF2* has been described in a patient displaying undescended testis^17^, possibly suggesting a role for *MCF2* in cellular migration during organogenesis.

The use of animal models is a powerful tool in order to mechanistically understand neurodevelopmental disorders^18^ in the context of rare genetic variants associated with complex traits^19^. The *in vivo* model used in the present study allowed us to identify an interesting pathophysiological process caused by the acute KD of *Mcf2* in the developing mouse. Further studies could take advantage of induced pluripotent stem cells and human organoids to recapitulate human developmental processes. Genomic analyses should be used to search for other rare variants associated with CBPS in a larger sample size to determine whether additional mutations in the MCF2 gene can be identified in rare cases of CBPS with similar lower motor neuron dysfunction. The convergence of additional genetic associations and *in vivo/in vitro* causal studies would allow to strengthen the pathophysiological mechanism identified in this study.

## Supplementary materials and methods

### q-RT-PCR

HEK 293T cells were cultured and transfected as previously ^8^ with the following plasmids: pUB6-m*MCF2* and *Mcf2* shRNA or *Scram* shRNA RNA was extracted from the sorted cells using a ReliaPrep™ RNA Cell Miniprep kit (Promega). 2 ng of RNA per sample was used for cDNA preparation, and pre-amplification and Real-time quantitative PCR was performed as previously described ^7^, using the following primers: *Mcf2* (forward: CCCTCAAAATTTCCCTCCAGA; reverse TGTAAGCTCCCAGGGGAAGGA) *ActinB*(forward: CTAAGGCCAACCGTGAAAAGAT; reverse: CACAGCCTGGATGGCTACGT), *Gapdh* (forward: TCCATGACAACTTTGGCATTG; reverse: CAGTCTTCTGGGTGGCAGTGA), *Tuba2* (forward: AGGAGCTGGCAAGCATGTG; reverse: CGGTGCGAACTTCATCGAT).

### Tissue processing and immunohistochemistry

P0 brains were dissected and soaked in 4% paraformaldehyde at 4°C overnight, cut in 50 um-thin coronal sections using a vibratome and stored at 4°C in 0.1M Phosphate Buffer Saline until further processing. Free-floating immunohistochemistry was performed as described^7^, using a mouse anti-HA (1/1000, Biolegend) primary antibody and a secondary donkey anti-mouse Alexa-488 antibody (1/500, Invitrogen).

### Statistics and analyses

All described image analyses were done using Fiji^20^, using the cell counter plugin. All statistical analyses were performed on Prism software. The sample sizes and relevant statistical tests are specified for each result in the results section.

## Acknowledgements and author’s contributions and funding

AMC designed and performed all the *in vivo* and *in vitro* experiments, including all the analyses and wrote the manuscript. AD designed the animal model experiments and wrote the manuscript. JF identified the case and conducted the clinical investigations. MG and SL performed the exome sequencing experiment and analyses. This work was supported by the Institute of Genetics and Genomics of Geneva (to AMC and AD) and by the NCCR Synapsy (51NF40-185897). The authors disclaim no conflict of interest. The authors would like to thank the patient and his parents, AMC would like to thank Dr Eleanor D’Ersu for proofreading of the translation of the clinical results.

